# Sensitive whole mount *in situ* localization of small RNAs in plants

**DOI:** 10.1101/037978

**Authors:** Mouli Ghosh Dastidar, Magdalena Mosiolek, Michael D. Nodine, Alexis Maizel

## Abstract

Small regulatory RNAs are pivotal regulators of gene expression and play important roles in many plant processes. Although our knowledge of their biogenesis and mode of action has significantly progressed, we comparatively still know little about their biological functions. In particular, knowledge about their spatiotemporal patterns of expression rely on either indirect detection by use of reporter constructs or labor-intensive direct detection by *in situ* hybridization on sectioned material. None of the current approaches allows for a systematic investigation of small RNAs expression patterns.Here, we present a method for the sensitive in situ detection of micro-and siRNAs in intact plant tissues that utilizes both double-labelled probes and a specific cross linker. We determined the expression patterns of several small RNAs in plant roots and embryos.

## Introduction

Small (20-25nt) RNAs (smRNAs) regulate gene expression in eukaryotes by guiding the transcriptional and post-transcriptional gene-silencing machinery by base pairing to their targets. In plants, smRNAs are involved in development, response to phytohormones and nutrients and ensure genome integrity by mediating the epigenetic regulation of mobile repetitive elements. (Jones-Rhoades *et al.*, 2006; Rubio-Somoza *et al.*, 2009; Sunkar *et al.*, 2007). Our understanding of smRNA biogenesis and action has made significant progresses in the recent years and modern sequencing technologies have tremendously expanded the repertoire of smRNAs. Yet in comparison, the study of their function is lagging behind. A key aspect of the functional characterization of smRNAs is a precise description of their spatial and temporal patterns of expression at the cellular scale. Published examples have revealed that in plants smRNAs are expressed in discrete, tissue or cell type-specific pattern (Juarez *et al.*, 2004; Nogueira *et al.*, 2009; Chitwood *et al.*, 2009; Wollmann *et al.*, 2010; Ori *et al.*, 2007; Cartolano *et al.*, 2007; Douglas *et al.*, 2010; Marin *et al.*, 2010; Carlsbecker *et al.*, 2010; Miyashima *et al.*, 2011; Yu *et al.*, 2015; Husbands *et al.*, 2015). The most common method to infer the expression patterns of smRNAs at the cellular level consists the use of transgenic reporter genes encoding detectable products like Beta-Glucuronidase *(uid* A), Green Fluorescent Protein *(GFP)* and Luciferase *(LUC)*, that are placed under the control of putative small RNA precursor promoters (Carlsbecker *et al.*, 2010; Yu *et al.*, 2015; Marin *et al.*, 2010; Nogueira *et al.*, 2009; Nogueira *et al.*, 2007; Chitwood *et al.*, 2009). Alternatively, ubiquitously expressed reporters containing small RNA binding sequence have been used to reveal the small RNA expression pattern as area of absence of reporter expression (Parizotto *et al.*, 2004; Nodine and Bartel, 2010; Marin *et al.*, 2010). However, because this method produces an absence of signal and is influenced by the efficacy of the small RNA, it can be difficult to interpret, and requires the time-consuming, and not always possible, generation of transgenic plants. Direct localization by *in situ* hybridization with specific anti-sense probes provides high resolution and does not rely on the establishment of such transgenic reporters. Original protocols employed digoxigenin labelled anti-sense RNA probes (Kidner and Martienssen, 2004; Ori *et al.*, 2007; Juarez *et al.*, 2004; Douglas *et al.*, 2010; Cartolano *et al.*, 2007; Nogueira *et al.*, 2009; Chitwood *et al.*, 2009) but were supplanted by use of locked nucleic acid (LNA) oligonucleotide probes that provide increased sensitivity and specificity (Kloosterman *et al.*, 2006; Wheeler *et al.*, 2007; Válóczi *et al.*, 2006). However these protocols suffer from two main disadvantages that preclude their widespread adoption. These methods rely on the use of embedded and sectioned material as a template for hybridization. The generation of these sections is a tedious and lengthy process and some tissues such as the root are particularly challenging to sectioning. Second, the current protocols have a limited sensitivity. Small RNAs are often not be very abundant and current protocols do not take into account the high volatility of small RNAs during the numerous washing steps of the protocols (Pena *et al.*, 2009). The conventional formaldehyde fixation of tissue used in such protocols has been shown to result in substantial loss of smRNAs which can be prevented by the usage of EDC (N-(3-Dimethylaminopropyl)-N'-ethylcarbodiimide hydrochloride) that covalently cross-links smRNA 5’ ends to the protein matrix (Pall *et al.*, 2007; Pena *et al.*, 2009). Here, we present an *in situ* hybridization protocol for smRNAs that enables their detection in non-sectioned plant tissues. Our protocol relies on the use of double-labelled LNA probes and a smRNA-specific post-fixation step using EDC for superior sensitivity. We have employed this method to document the expression pattern of seven micro-and short interfering RNAs in Arabidopsis roots and embryos. This approach is highly specific and allows for a semiquantitative assessment of small RNA abundance. The whole procedure can be preformed using liquid-handling robotic systems and is therefore amenable to high-throughput applications.

## Results & Discussion

We devised a protocol for the detection of small RNAs in whole mount Arabidopsis samples (Figure 1). After initial steps of fixation, dehydration, proteinase treatment and post-fixation, samples are optionally treated with EDC (N-(3-Dimethylaminopropyl)-N'-ethylcarbodiimide hydrochloride), a chemical reacting with free 5’ ends present in small RNAs and condensing them with amino groups in the protein matrix of the tissue (Pall *et al.*, 2007; Pena *et al.*, 2009). This step ensures that the small RNAs are not washed away during subsequent steps (Pena *et al.*, 2009). We used LNA-modified probes for hybridization with fixed samples. LNA oligos are high-affinity RNA analogs in which the furanose ring of selected base is locked by an O2',C4'-methylene bridge resulting in increased hybridization properties and specificity towards their targets. The probes were chemically modified on their 5’ and 3’ ends with a digoxigenin residue. The presence of two digoxigenin group enhances the sensitivity of the probe. Hybridizations were performed overnight at temperatures of 55°C or 65°C depending on the probe (Table 1). After washing and blocking, the hybridized probe was detected by incubating with an antibody against digoxigenin coupled to an alkaline phosphatase followed by its colorimetric detection with chromogenic substrate under the microscope. As a proof of principle, we applied the protocol to the detection of miR390 in the root system of 7-day old Arabidopsis wild type (Col-0). Mir390 is an abundant miRNA in the root produced by a single active locus *(MIR390a)* (Breakfield *et al.*, 2011; Marin *et al.*, 2010), and expressed in the meristem region of the primary and lateral root primordia (Marin *et al.*, 2010). The specificity of the detection was assessed by performing the procedure either with a probe antisense to the miR390 sequence, a sense probe or no probe at all. We observed strong hybridization signals in the primary root, as well as in the lateral root primordia with the antisense miR390 probe (Figure 2A). We observed very weak to no signal in samples incubated with either the sense or no probe at all (Figure 2B, C). In the samples hybridized with the miR390 antisense probe, signal was more prominent in the cells of the meristem regions in particular in the stele, whereas signal in the endodermis, cortex and epidermis were also observed. No signal was observed in the columella cells. The miR390 pattern observed by *in situ* hybridization was identical to that produced by the transcriptional reporter pMIR390a::GUS:GFP (Marin *et al.*, 2010). MiR390 signal was only detected in the cytoplasm of the cells and more easily visible in cells without a fully developed vacuole (Figure 2D). Together, these results indicate that miR390 can be specifically detected in whole mount Arabidopsis root samples.

**Table 1.**
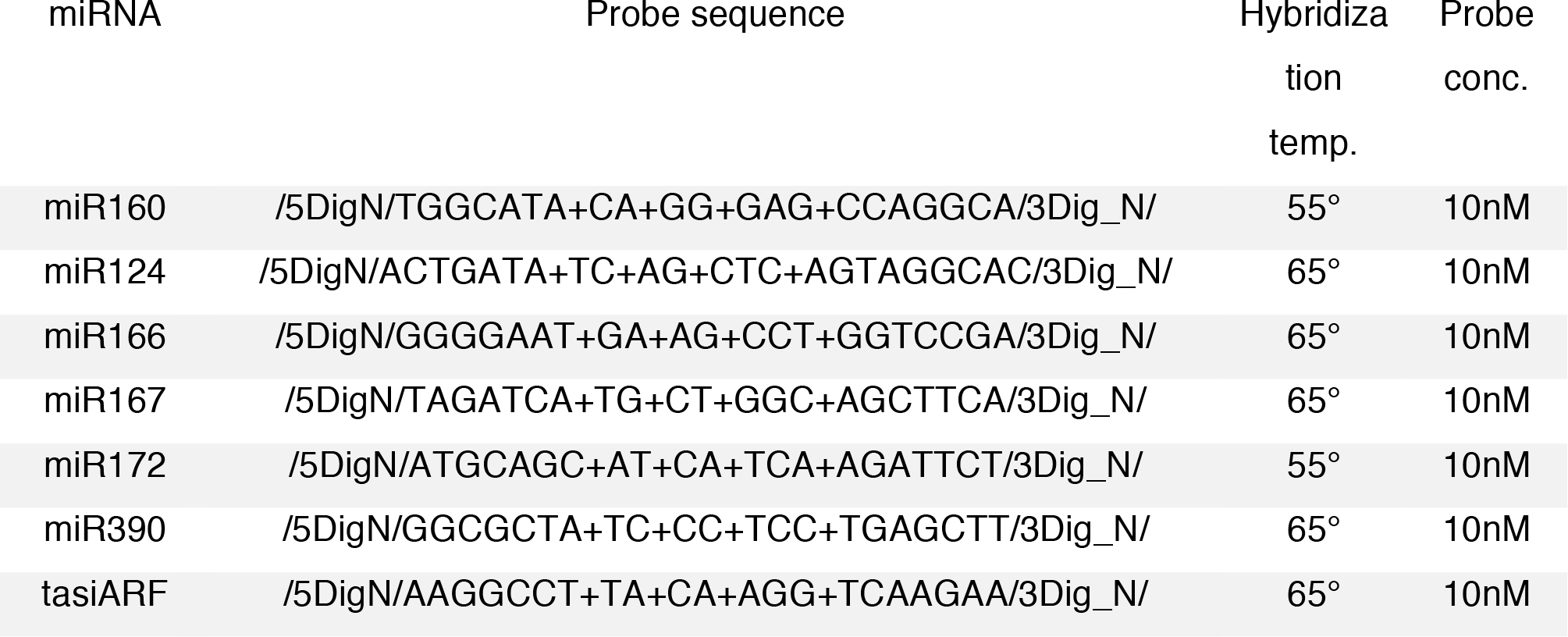
Different microRNAs and their probe sequences, hybridization temperatures and probe concentrations

**Figure 1.**
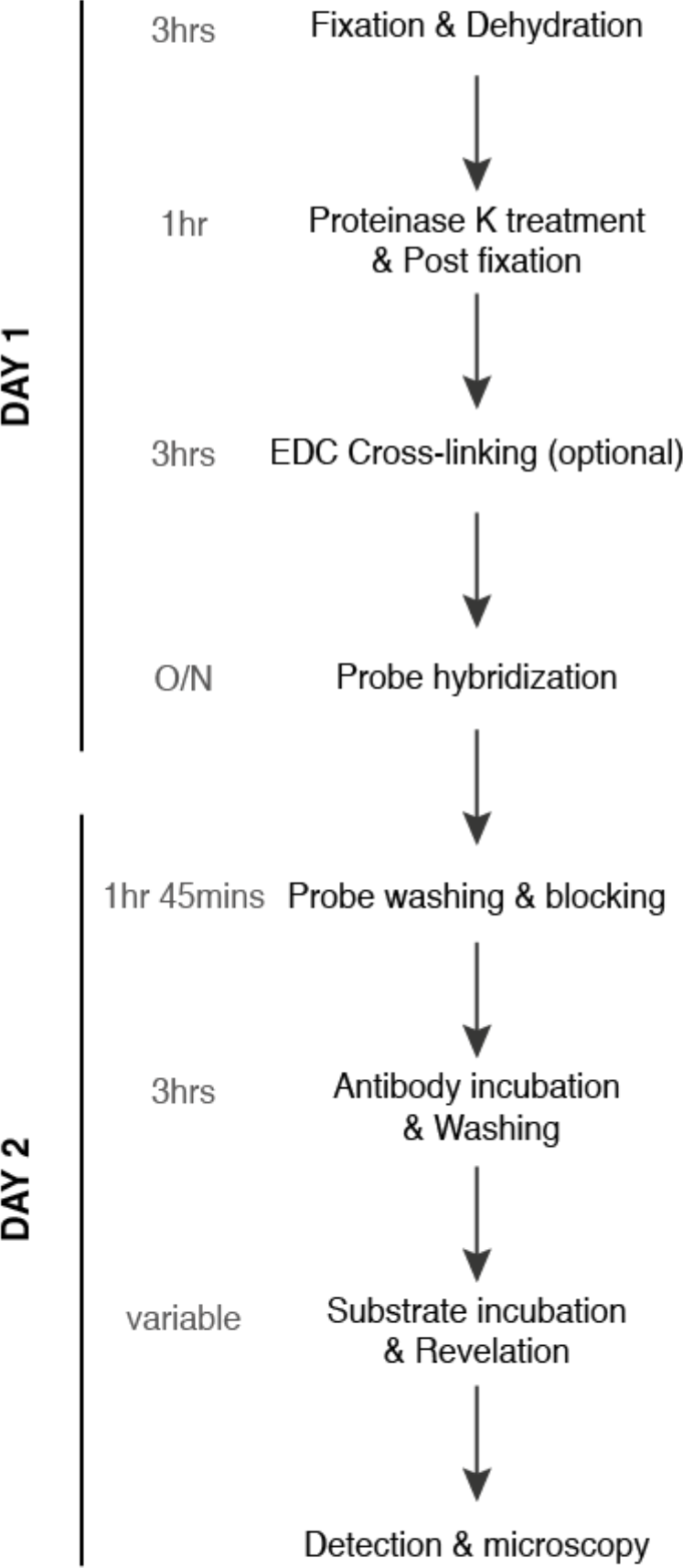
A detailed step-by-step description of the procedure can be found in the supplemental material.

**Figure 2.**
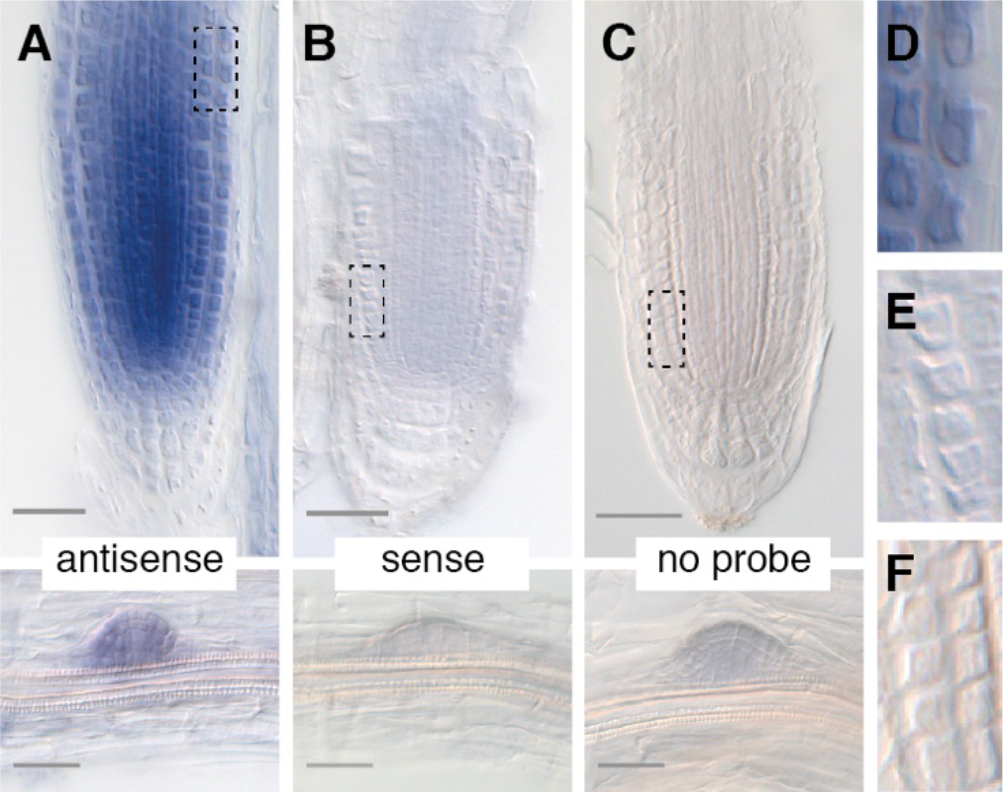
Visualization of miRNAs in whole mount Arabidopsis roots. Whole mount *in situ* hybridization of double digoxigenin labelled locked nucleic acid miRNAs probes carried out on *Arabidopsis thaliana* (wild type, Col-0) roots. miR390 signals were specifically detected both in primary root (upper panels) and lateral root primordia (lower panel) only with the antisense probe (A). Little to no signal were detected with the sense (B) or in absence of probe (C). (D-F) Higher magnification views of the areas delineated in (A-C). Scale bars are 50*μ*m.

We then tested other less abundant root miRNAs (Breakfield *et al.*, 2011) such as miR160, miR166 and miR167. Whereas miR390 could be easily detected in absence of EDC-crosslinking, in the same conditions, we did not detect any signal for samples incubated with miR160, 166 or 167 antisense probes (Figure 3A, C, E). In samples treated by EDC, we observed signal in the primary root meristem region with all three probes (Figure 3B, D, F). Similar to miR390, these additional miRNA signals were cytoplasmic and easier to visualize in non-fully vacuolated cells. The signal obtained with the miR166 antisense probe was more intense in the epidermis and cortex layers than in the inner layers (Figure 3D), which is in agreement with previous reports (Carlsbecker *et al.*, 2010). Whereas miR160 signal was also more intense in the outer layers (Figure 3B), miR167 was more intense in the stele (Figure 3F). miR172, did not yield any strong signal even in presence of EDC (Figure 3G, H), although it appears to be expressed in roots (Breakfield *et al.*, 2011). We verified that the EDC treatment preserved the specificity of the detection by probing Arabidopsis roots treated with EDC or not with probes against the animal-specific miR124. We did not detect any signal even when the samples had been cross-linked by EDC (Figure 3I, J). In their ensemble, these results show that EDC-crosslinking of small RNAs improves the sensitivity of detection while preserving the specificity.

**Figure 3.**
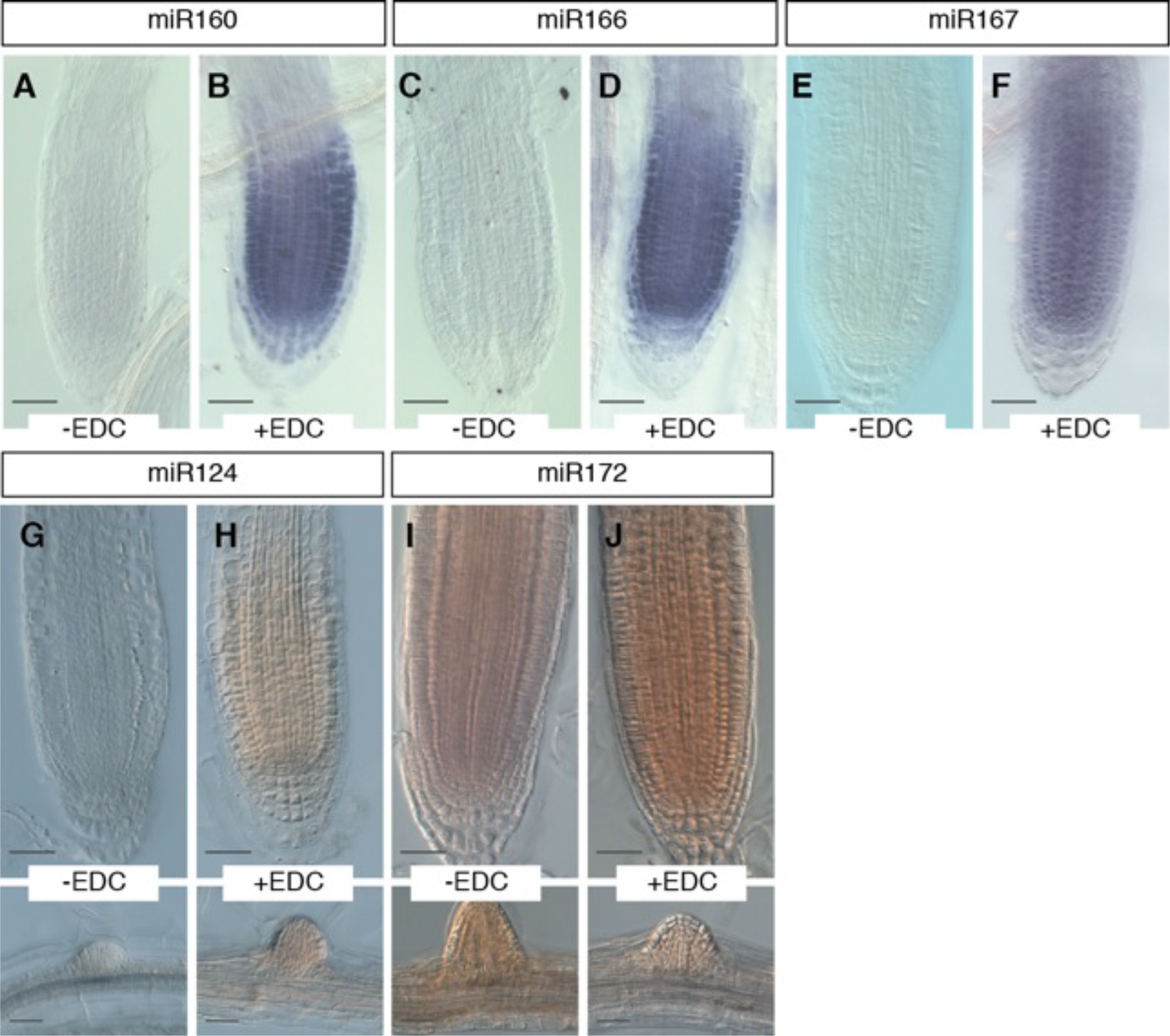
Increased sensitivity by use of EDC fixation. Expression of miR160, miR166, miR167, miR124 and miR172 in absence (A, C, E, G, I) and in presence of EDC crosslinking (B, D, F, H, J). EDC-treatment allows the detection of miR160, 166 and 167 signal. In the same conditions, the mouse miRNA miR124 and the Arabidopsis miR172 do not show any signal. Scale bars are 50*μ*m.

To extend our observations to other tissues, we applied our smRNA *in situ* method to Arabidopsis embryos with a few modifications as outlined in the experimental procedures. Signals corresponding to miR160, miR166, miR167 and miR390, but not miR172, were reproducibly detected above the background signal observed when using the animal-specific miR124 as a negative control (Figure 4). Consistent with the results from post-embryonic roots, we also observed the localization of miR160, miR166, miR167 and miR390 in embryonic radicles. In addition, miR160, miR166 and miR167 were also present in the shoot meristem region and the hypocotyl of embryos. Similar to the results from post-embryonic roots, the use of EDC-crosslinking in the WISH procedure improved the sensitivity of miR160, miR166 and miR167 although this was not required to detect miR390. Moreover, because miR160 probes give strong signal in the innermost cell layers (i.e. vascular tissue precursors) in either the presence or absence of EDC, the use of EDC-crosslinking preserved the cell-specificity of the *in situ* hybridizations (Figure 4A-B).

**Figure 4.**
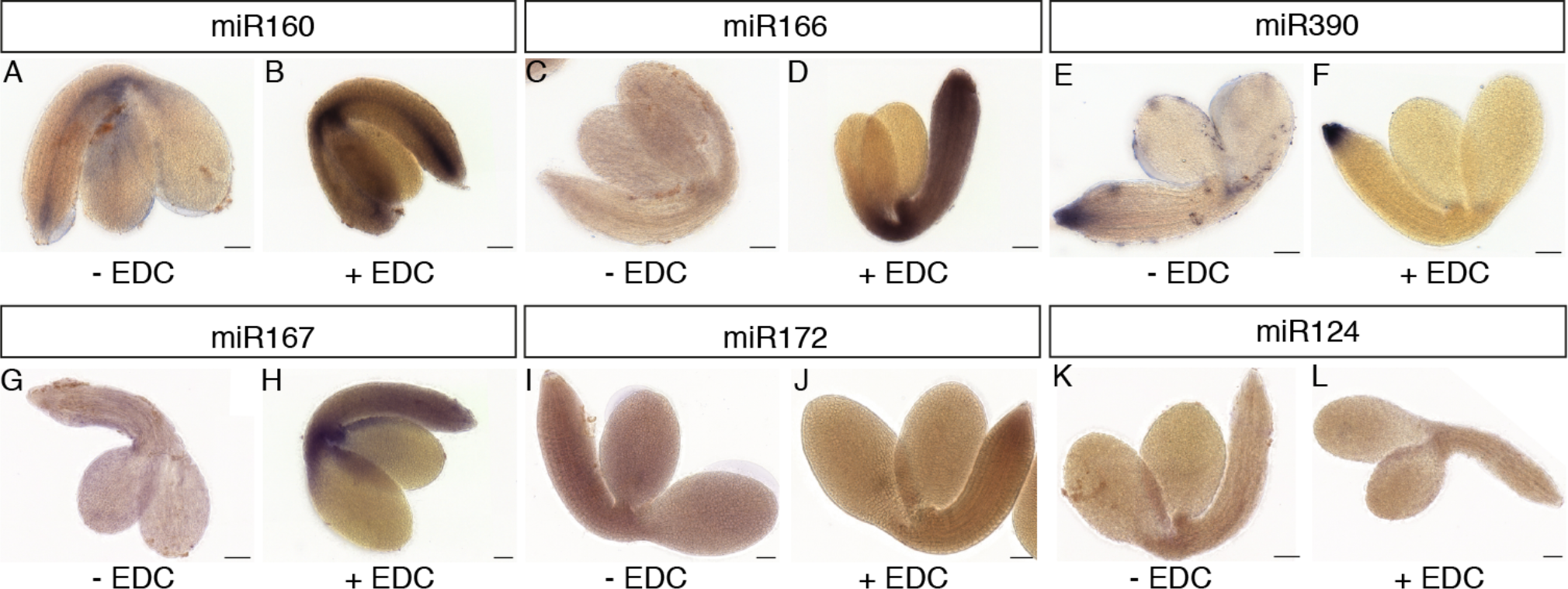
miRNA detection in embryos. Images of embryos hybridized with probes corresponding to miR160, miR166, miR390, miR167 and miR172 *in situ* hybridizations in either the absence (A, C, E, G, I) or presence of EDC-crosslinking (B, D, F, H, J) are shown. Probes against the mouse miRNA miR124 were used as negative controls and were performed either without (K) or with (L) EDC-crosslinking. Scale bars are 50*μ*m.

Given the sensitivity of the method, we tested whether it would be amenable to the qualitative assessment of the abundance of other small RNAs. For this purpose, we designed a probe against the trans acting (ta)-siRNAs produced by the *TAS3* pathway (tasi-ARFs) and observed tasi-ARF accumulation in either wild typeor in lines in whichthe expression of the *TAS3a* precursor iseither reduced *(tas3a-1*, (Adenot *et al.*, 2006)) or it is over-expressed *(TAS3 OX*, (Marin *et al.*, 2010)). *TAS3 OX* plants have a 2-3 fold increased in tasi-ARFs levels compared to wild type (Marin *et al.*, 2010) whereas *tas3a-1* mutant has only 40% of wild-type levels (Adenot *et al.*, 2006). All samples were treated in parallel and the incubation with the chromogenic substrate was identical for all three genotypes. The hybridization signal intensities observed were lower in the primary of *tas3a-1* mutants compared to wild type (Figure 5A, C) and stronger in the *TAS3 OX* overexpression lines (Figure 5B). These results demonstrate that the method is able to detect siRNAs and can pick up relative differences in their abundance.

**Figure 5.**
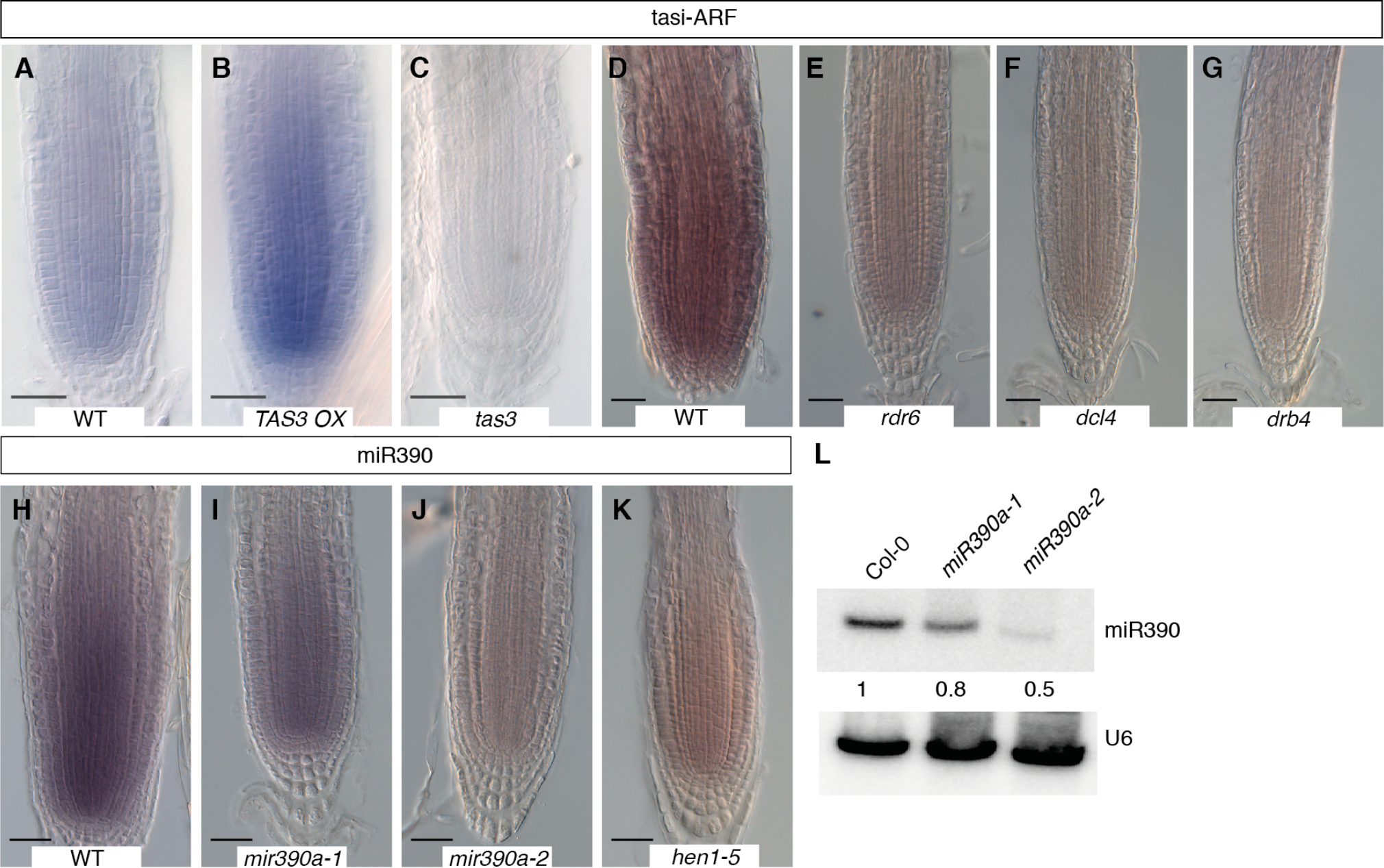
Semi-quantitative detection of small RNA abundance. (A-G) Detection of TAS3-derived ta-siARFs ta-siRNAs in roots of wild type plants (A, D), in plants expressing the precursor *TAS3* (B) or in plants impaired in ta-siRNAs biogenesis such as the *tas3* mutant (C), the *rdr6* (E), *dcl4* (F) or *drb4* (G) mutants. (H-K) Detection of miR390 in wild type (H) plants, a in mutant partially affected in miR390 processing from the *MIR390a* precursor *(mir390a-1*, I), in a mutant with reduced levels of the precursor (*mir390a-2*, J), in a mutant destabilizing mature miR390 by lack on terminal methylation (*hen1-5*, K). Signals are reduced in the *miR390a-1* mutant and absent in *mir390a-2* and *hen5* mutants. Scale bars are 50*μ*m. (L) Northern blot for miR390 in root RNA of 7-day old wild type, *mir390a-1* and *miR390a-2* seedlings confirming the reduction of miR390 abundance in these mutants. U6 serves as loading control. Values indicate relative abundance of miR390 compared to wild type (Col-0).

To further validate the applicability and specificity of the method, we probed for abundance of small RNAs in plants impaired in the biogenesis of these small RNAs. We first tested the abundance of tasi-ARFs in *rdr6, dcl-4* or drb4mutant plants that are deficient in tasi-ARF production (Elmayan *et al.*, 1998; Gasciolli *et al.*, 2005; Adenot *et al.*, 2006; Xie *et al.*, 2005). Whereas signal was detected in wild type, no signal could be seen in the *rdr6, dcl-4* or drb4mutants (Figure 5E, F, G). We then compared the levels of miR390 in wild type and inthe *miR390a-1, miR390a-2* and *henl-5* mutants. Whereas *miR390a-1is* a mutant defective in the processing of the MIR390a precursor (Cuperus *et al.*, 2009), *miR390a-2has* reduced levels of the MIR390A precursor (Marin *et al.*, 2010). *Thehen1-5* is a mutant impaired in miRNA biogenesis(Park *et al.*, 2002). The signal intensities were lower in *miR390a-1* mutants compared to wild type and even lower in *miR390a-2* (Figure 5H, I, J) and almost no signal could be detected in *hen1-5* mutant (Figure 5K). The reduced levels of miR390 in the *miR390a-1* and *miR390a-2* mutants was validated by northern blot analysis, which showed an 80% and 50% reduction of mature miR390 levels in respective *miR390a-1* and *miR390a-2* mutants compared to wild type (Figure 5L).

In summary, this protocol is sensitive, robust and efficient as it does not involve the long steps of tissues embedding and sectioning before hybridization. The use of double labelled LNA probe and the stringent hybridation conditions ensure both sensitive and specificity to the method as exemplified by the very low of absence of signal when using either sense probes, unrelatedmiRNA (miR124) or performing hybridization in mutant backgroundswith reduced miRNA abundance. Moreover, the protocol is applicable to both embryonic and mature tissues and amenable to the detection of miRNAs and siRNA (ta-siRNA). The protocol should be applicable to other tissues or plant species. The step of small RNA cross-linking with EDC improves the detection of several small RNAs while preserving specificity. Here we have used an additional step of cross-linking of small RNAs by EDC (N-(3-Dimethylaminopropyl)-N'-ethylcarbodiimide hydrochloride) which improved the sensitivity of detection in particular for low abundant small RNAs. Signals for three miRNAs in particular-miR160, miR166, miR167 could be enhanced specifically in the primary roots in contrast to non-EDC treated roots where levels of miRNAs were low or below detection level. The method allows the detection of relative differences in the abundance of small RNAs or to profile small RNAs in mutant backgrounds. Theexpression patterns obtained for the different miRNAs corroborated the ones inferred using indirect methods (miR390, miR166, miR167, miR160 (Marin *et al.*, 2010; Breakfield *et al.*, 2011; Carlsbecker *et al.*, 2010)). In addition, the comparison of the expression patterns between the embryo and the seedling revealed the dynamics of miR172 expression, detected in embryo but less abundant in young plants.Importantly, the protocol is amenable to high-throughput applications since the whole procedure can be performed using liquid-handling robotic systems. This improved method allows the systematic determination of a wide range of small RNAs localization patterns and subsequent inference of their functions.

## Experimental procedures

### Plant material

All lines used in this study are in the *Arabidopsis thaliana* Col-0 ecotype background. All mutant used have been previously described: *mir390a-1* (Cuperus *et al.*, 2009), *mir390a-2* (Marin *et al., 2010),rdr6* (sgs2-1)(Elmayan *et al.*, 1998; Mourrain *et al.*, 2000), *dcl4-1(Xie et al.*, 2005), *tas3a-1, OXTAS3* (Marin *et al.*, 2010), *drb4* (Adenot *et al.*, 2006), *hen1-5* (Park *et al.*, 2002). Embryos were dissected in a drop of PBS and immediately transferred to 4% paraformaldehyde on ice.

### Plant growth conditions

Plants were grown on 0.5X-MS/0.8% agar (MS-agar) plates in controlled-environment chambers under the following conditions: 150 *μ*mol photon.m^-2^.S^-1^ luminance, 16 hr light, 23°C temperature.

### Probe design

LNA-modified oligonucleotide probes were synthesized by Exiqon. The probes corresponds to the full antisense sequence of the miRNA, are DIG-labeled at both the 5’ and 3’ ends and contain 4 LNA-modified base at positions 7, 9, 11, 15 (Table 1).

### Whole mount in situ hybridization

Only the main steps of the method are described here, a detailed protocol is available as supplemental material. Tissues were cross-linked with para-formaldehyde (4% solution) by vacuum infiltration until the samples sunk and became translucent in color. Length of fixation varied according to the tissue. Root and embryo samples are typically fixed for 45 minutes and 2 hours at room temperature, respectively. Once fixed, samples are permeabilized and chlorophyll was removed by a 1:1 mixture of 100% ethanol and Histo-Clear®. Cleared root tissues and embryos were then digested by either 125 *μ*g/mL proteinase K for 30 minutes or 75 *μ*g/mL proteinase K for 15 minutes, respectively. For the optional EDC (N-(3-Dimethylaminopropyl)-N'-ethylcarbodiimide hydrochloride) cross-linking step, incubation was done at 60°C for 2hrs at 0.16M concentration. The optimal hybridization conditions (temperature and probe concentration) were optimized for each probe. For seedlings, probes were typically hybridized at 55°C or 65°C at 10nM for 16h. For embryos, all probes were hybridized at 65°C with 20nM of probe for 16h.After hybridization, samples were washed and incubated with Digoxigenin antibodies followed by NBT/BCIP incubation in the dark for colorimetric detection. The time of revelation depends on the probe. For root and embryo samples, revelation was done for 1 hour (miR390), 2-4 hours (miR166, miR167 and miR172), 3 hrs for ta-siRNA and 6 hours (miR160). As negative controls, *in situ* hybridizations using animal-specific miR124 probes were performed on root and embryos alongside and under the exact same conditions as the miRNAs tested.

### Imaging

For imaging roots, samples were mounted in a 1:1 mixture of glycerol and 1X TE buffer and detected by light microscopy using Nomarski (Differential Interference Contrast, DIC) from Nikon with a 20X objective. After stopping the colorimetric detection, embryos were transferred to three-welled glass slides (Electron Microscopy Science Cat. No. 63418-11), mounted in 70% glycerol/TE buffer and sealed with cover-slips. Slides were subsequently imaged on an automated Pannoramic SCAN 150 slide scanner (3DHISTECH) with transmitted light and a 20X plan-apochromat objective. Images of embryos were collected using the Panoramic Viewer software (3DHISTECH).

### Northern blotting

Northern blotting was performed as described (Marin *et al.*, 2010)

## Acknowledgements

We thank A. Bleckmann for her comments on the manuscript. This work was supported by the Land Baden-Württemberg, the Chica und Heinz Schaller Stiftung, the CellNetworks cluster of excellence and the Boehringer Ingelheim (to M.G.D, A.M.), and the framework of the first call ERA-NET for Coordinating Plant Sciences (ERA-CAPS) funded by the European Union in the framework of the FP7 under the Grant Agreement n°291864, with funding from the Austrian Science Fund (FWF grant number 1476-B16) (M.M and M.D.N).

## Supporting Information

Additional Supporting Information may be found in the online version of this article.

